# Lizard colour plasticity tracks background seasonal changes

**DOI:** 10.1101/862268

**Authors:** Daniele Pellitteri-Rosa, Andrea Gazzola, Simone Todisco, Fabio Mastropasqua, Cristiano Liuzzi

**Author notes:** Author for correspondence: Daniele Pellitteri-Rosa.

## Abstract

Environmental heterogeneity on spatial and temporal scale fosters organism’s capacity to plastically alter coloration. Predation risk might favour the evolution of phenotypic plasticity in colour patterns, as individuals, which change colour throughout the year, could be able to improve their fitness. Here we explored the change in dorsal pigmentation of the Italian wall lizard (*Podarcis siculus campestris*) along three time points (March, July and October) during the period of activity. Lizard dorsal pictures were collected on the field, with the support of a reference chart to quantitatively estimate chromatic variables (hue, saturation and value, HSV). At the same time, pictures of grassy coverings (the most representative portion of the environment subjected to normal seasonal change), were collected. Our findings show that lizards are capable of altering dorsal coloration during seasonal change. They vary from green, at the onset of spring, to brownish in the middle of summer, and greyish colour in October. This modification closely followed environmental background colour variation and enhanced lizard crypsis during each season.

## 1. Introduction

Differences in cryptic coloration among animals have provided evidence for significant selective advantages in specific contexts [1,2,3]. One possible trigger of chromatic variation is the habitat-specific background colour. When the level of predation is high, a body coloration matching the background colour is common among prey species and has been reported in many studies [4,5]. The visual resemblance to the background chromatic characteristics (colour, brightness, or pattern) has been proved to be an adaptive trait, which can significantly decrease the risk of detection by predators [6]. Natural populations provide examples of habitat-specific body colours in a great diversity of taxa, including freshwater fish [7], frogs [8], salamanders [9], turtles [10], lizards [11], and mice [2]. Such studies suggest that the adjustment to habitat-specific variation in colour is mostly a result of selective pressure by predators hunting by sight, leading the population towards an adaptive match to the local background. However, body coloration can also affect the fitness of animals through its effects on thermoregulation [12] or social status [3].

Animal colour regulation may be fixed (i.e. constitutive), for the past effect of natural selection, or expressed over different timescales (from seconds to years) and is regarded as a form of phenotypic plasticity. In order to achieve an effective background matching, colour change can also follow seasonal changes. This phenomenon has been observed in numerous invertebrates and vertebrates. The tropical butterfly *Bicyclus anynana* shows seasonal variation in wing patterns, following the temperature which anticipates seasonal change in vegetation [13]; the snowshoe hare (*Lepus americanus*) goes through seasonal coat colour shifts from brown to white. This change has been recorded to be strongly related to survival [14].

Lizard coloration has been explored throughout numerous ecological and evolutionary studies [15,16,17], however seasonal colour change has been rarely explored in this group [18,19]. In this study, dorsal colour variation in *Podarcis siculus campestris* was recorded by sampling individuals of the same population in three different months during the yearly period of activity. We also explored the possible differences between sexes in colour variation, and whether it might enhance concealment by tracking environmental modifications which occur during seasonal changes.

## 2. Material and Methods

### (a) Animal sampling

During 2018, we conducted field preliminary observations of adult lizards in a population from central Apulia (southern Italy) throughout their period of activity. This was done by collecting dorsal pictures (Sony HX300 resolution 5184 × 3888 pixels) without capturing the lizards. A first rough visual inspection of the pictures and the number of photographically recaptured lizards (59 out of 356 total observations) allowed us to assess an individual seasonal colour variation (see electronic supplementary material, S1, for additional details). In 2019 we collected adult lizards in the same population during three capture sessions temporally distributed as follows: March (late winter, early spring), July (full summer) and October (late summer, early autumn). The sampling site is included in the European special area of conservation (SAC) “Murgia dei Trulli”, whose landscape is singularly characterized by typical dry constructions with conical roofs (“trulli”), surrounded by oak trees, olive groves and a large covering of grassy vegetation. The animals were captured by noosing and individually kept in cloth bags until the end of sampling. The lizards were then sexed and measured using a digital calliper (accuracy ± 0.1 mm) for the snout-vent length (SVL). As for many lizard species, males are clearly distinguished by females both in their larger body and head dimension and in their well-developed femoral pores [20]. The belly was photographed to check for possible recaptures over the sessions by means of the I^3^S software on the ventral scales [21] (see electronic supplementary material, S2, for additional details). In order to measure the colour variation through the seasons, we took a picture of the dorsal body region of each lizard adjacent to a GretagMacBeth Mini Color Checker chart (24 colour references, 5.7 cm × 8.25 cm). During each sampling session, we also took 20 pictures of grassy vegetation adjacent to the Mini Color Checker chart (see electronic supplementary material, S3, for additional details).

This was carried out to quantitatively characterize the vegetation cover colour representing the entire area considered in the sampling. The lizards were then released at the exact site of capture. Overall, we captured 135 lizards (80 males - 55 females, mean SVL ± se = 74.88 ± 0.54 mm and 63.37 ± 0.65 mm respectively), well distributed over the seasons (March: 28 - 18; July: 27 - 21; October: 25 - 16).

### (b) Colour measurement

We used RGB values both for lizards and the environment by adopting the method initially proposed in monkeys [22] and later also used in reptiles [23]. We used the Camera plug-in for Adobe Photoshop CS6 to create a new colour profile that adjusted the colour in the photographs (in the tiff format) to the known colour levels in each square of the ColorChecker chart. For each lizard, we measured the colour of the dorsal part by selecting the areas of all scales showing colouration (i.e. black spots were excluded), using the ‘magic wand’ tool (on average roughly 93,000 pixels) and recording the RGB levels using the histogram palette. The RGB colour values were then rearranged in the Hue, Saturation and Value (HSV) system which is the most used representation of points in an RGB colour model.

### (c) Statistical analyses

With the aim to explore the seasonal change in lizard dorsal colour, we adopted linear mixed models which included SVL as covariate, and sex, month and their 2-way interaction as fixed factors. The date of data collection was included as a random factor. We ran three models with hue, saturation and value as response variables, respectively. In order to meet the assumption of residuals homogeneity, we included a variance structure for a different spread per stratum (i.e. month). This resulted in a better fit and lower AIC of the model. In all the models, we assumed a normal error distribution. Comparisons between lizard and background colour for each month were obtained through a series of Wilcoxon rank sum tests for hue, saturation and value, respectively.

## 3. Results

Hue varied as function of both month and sex (month × sex, *χ*^*2*^ = 18.23, *df* = 2, *p* <0.001), showing the highest value during March and decreasing in July and October (Fig.1), but was not affected by SVL (*χ*^*2*^ = 0.11, *df* = 1, *p* = 0.73). Saturation showed a significant effect of month (*χ*^*2*^ = 45.96, *df* = 2, *p* < 0.001) but not of sex (*χ*^*2*^ = 1.51, *df* = 1, *p* = 0.21) and month × sex interaction (*χ*^*2*^ = 4.20, *df* = 2, *p* = 0.12), but SVL resulted significant (*χ*^*2*^ = 6.01, *df* = 1, *p* = 0.01) and was related to a proportional increase of saturation (slope ± s.e. = 0.21 ± 0.09, *df* = 123, *t* = 2.47, *p* = 0.015; Fig.1). Mixed model for value showed a non-significant effect for month, sex and their interaction (*χ*^*2*^ = 1.84, *df* = 2, *p* = 0.39; *χ*^*2*^ = 0.01, *df* = 1, *p* = 0.92; *χ*^*2*^ = 2.64, *df* = 2, *p* = 0.26, respectively); SVL was not significant (*χ*^*2*^ = 2.77, *df* = 1, *p* = 0.1) and showed a negative relationship with value (slope ± s.e. = −0.17 ± 0.10, *df* = 123, *t* = −1.66, *p* = 0.09; Fig. 1).

**Figure 1.**
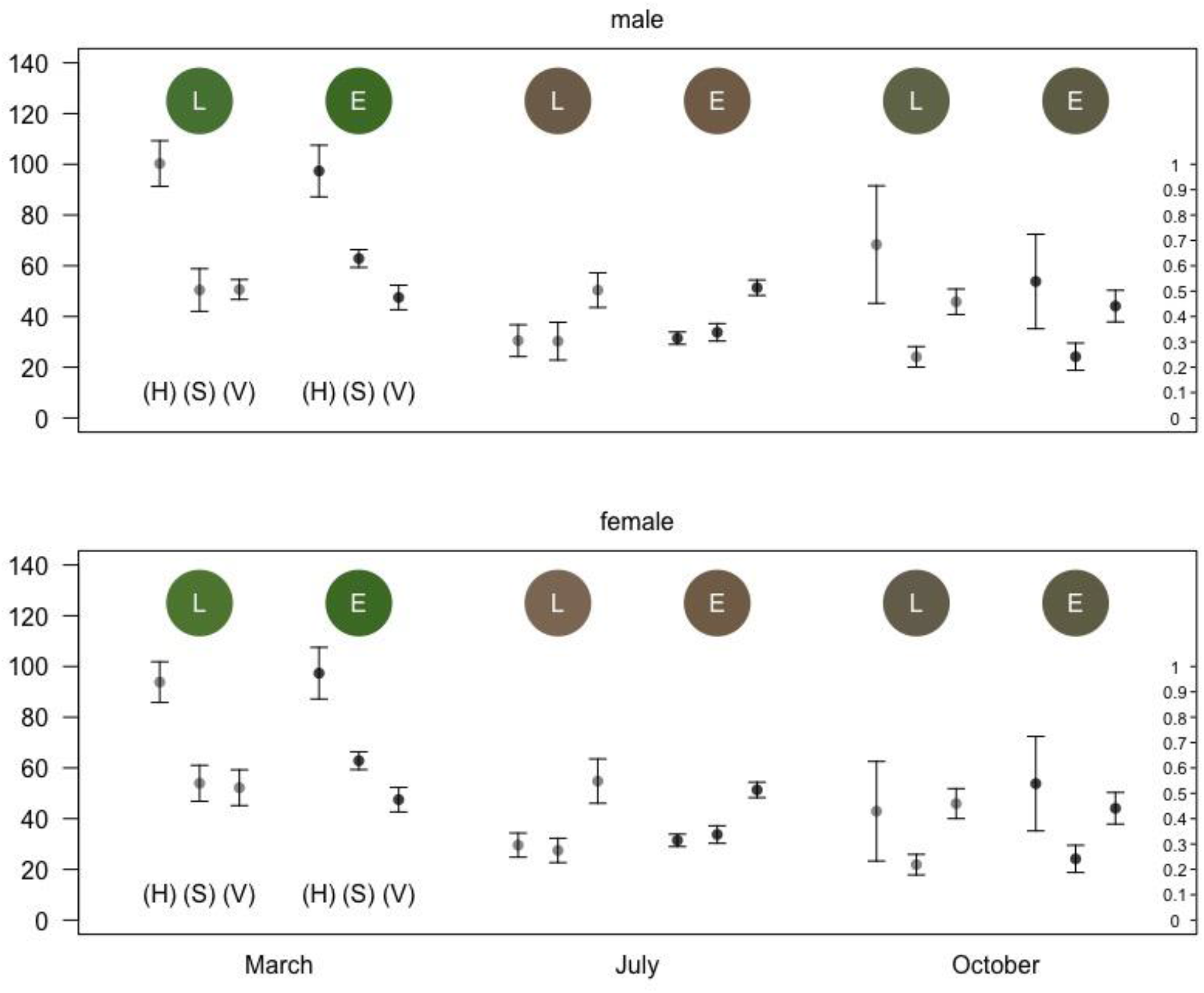
Means ± standard deviations of hue (H), saturation (S) and value (V) for both lizards (light grey) and grassy habitat (dark grey) in the three months of data collection on the field (March, n = 46; July, n = 48; October, n = 41) during 2019. For each plot (males and females), the colours of both lizard (L; left top coloured circular patch) and environment (E; right top coloured circular patch), generated by means values of HSV, are reported. The y-scale on the left is referred to H (°), the one on the right is related to S and V (%).

In March, the lizard hue matched the hue of the environment (*W* = 450, *p* = 0.89) but saturation and value did not (*W* = 77, *p* < 0.001 and *W* = 631.5, *p* = 0.01, respectively). In July, the lizard mean saturation estimate was different in comparison to the saturation of the environment (*W* = 244.5, *p* = 0.001), but hue and value were similar (*W* = 411.5, *p* = 0.35, *W* = 558, *p* = 0.29, respectively). To conclude, all hue, saturation and value matched the environment in October (lowest *p* = 0.53).

## 4. Discussion

In this study we detected an evident seasonal dorsal colour variation in the Italian wall lizard. Our findings reveal that at the start of spring, animals show a typical green colouration with greater hue values with respect to both those saturation and lightness; during summer, lizards exhibit a less lively dorsal colour tending to brownish, with a strong decrease of the average hue; finally, at the beginning of autumn, dorsal colouration shifts to greyish with an increase in hue and a reduction in saturation, which corresponds to a grouping of both green and brownish lizards. Moreover, through image analyses and photographic identification, we verified that colour varies on an individual basis and, therefore, not only at the population level. To our knowledge, studies on seasonal colour variation in lizards are few and related more on sexual selection or thermoregulation in controlled experiment contexts [18; 19]. Our results should be taken into account since many studies have described lizard taxonomic units based on chromatic patterns of a single season [24]. Systematic studies should probably be reconsidered, in view of possible colour changes throughout different seasons.

In the Italian wall lizard, the observed sharp change in dorsal colouration throughout the seasons could be associated with an anti-predatory adaptation. This is done by displaying a colouration resembling that of those backgrounds in which they are the most exposed to potential predators [25]. Interestingly, we found a similar seasonal trend in background mean values of hue, thus revealing an environmental colour matching with lizard dorsal tonality. The grassy habitat, which in our study area is the main microhabitat used by lizards for thermoregulation, radically changes between spring and summer. The landscape goes from a bright and intense green in spring to a brownish colour in summer due to the dry and arid grass (Fig. 2). In late summer, with the onset of the first rains, the grass begins to regain a greenish colour, creating a mosaic of colours ranging from brown to green, giving an overall shade of grey to the environment. Mean values of chromatic variables for both the lizards and the grassy habitat clearly show that lizard colouration strongly matched seasonal changes of the environment, providing arguments for adaptive cryptic adjustment [26] (see electronic supplementary material, S4, for additional details). The characteristic Mediterranean area considered in this study included also stone walls, potentially useful for lizard thermoregulation, though less frequently used by them. In contexts where a potential prey moves on different backgrounds, a camouflage strategy can be adopted only on a singular background, for example, the most frequent in the environment or where they are more visible to predators [27].

**Figure 2.**
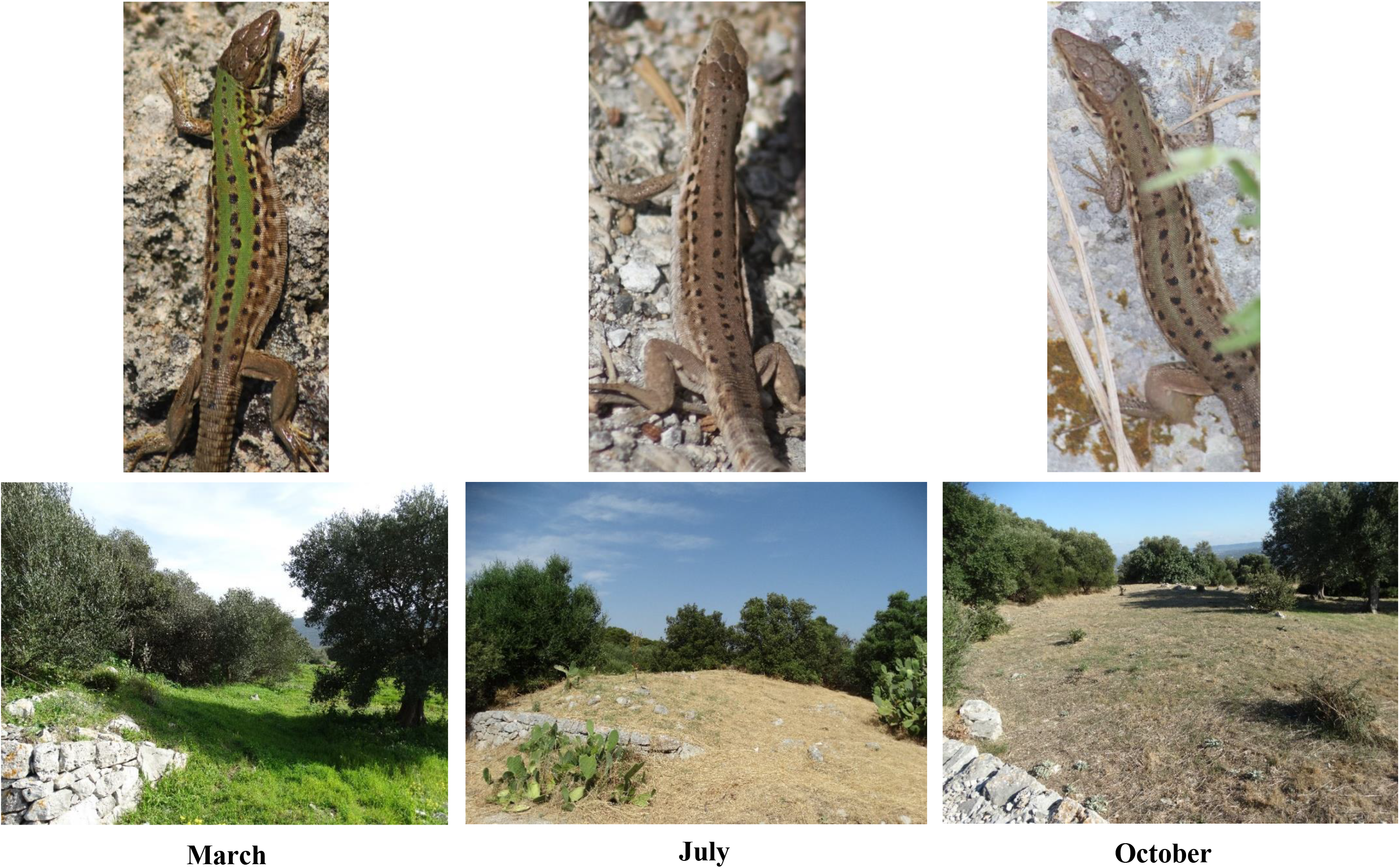
Matching of colour variation of lizards and grassy habitat throughout the different seasons. The lizard in the picture is the same female individual photographically recaptured in all the three sampling sessions.

We also found interesting results concerning both sex and size/age in relation to colour variation. Males and females did not show particular differences in chromatic variables throughout the sessions. This suggests that dorsal colour variation may be a generalized phenomenon, without implications related to differential sex-dependent strategies. However, saturation and value appeared to be affected by size, thus indicating a possible effect of the individual’s age on colour expression, as known for other lizard species [28,29].

## Ethics

All lizards captured in this study were kept in cloth bags during one-day sessions and later released at the exact site of capture, thus minimizing the disturbance to their biorhythms. This study was realized in conformity with the current Italian laws (DPN/II DIV/45377/PNM/2019).

## Data accessibility

The dataset is available from the electronic supplementary material, S3.

## Authors’ contributions

CL and DP-R developed the study concept; CL, DP-R, ST and FM contributed to the animal sampling and picture collection; DP-R and AG examined digital images and ran the statistical analyses; DP-R and AG drafted the manuscript. All authors have contributed to the revisions of this manuscript, agreed to be held accountable for this work and approved the final version of the manuscript for publication.

## References

1. Endler JA. 1981 An overview of the relationships between mimicry and crypsis. Biol. J. Linn. Soc. 16, 25–21.

2. Hoekstra HE. 2006 Genetics, development and evolution of adaptive pigmentation in vertebrates. Heredity 97, 222–234.

3. Stuart-Fox D, Moussalli A. 2009 Camouflage, communication and thermoregulation: lessons from colour changing organisms. Philos. Trans. R. Soc. B 364, 463–470.

4. Endler JA. 1978 A predator’s view of animal color patterns. Evol. Biol. 11, 319–364.

5. Ruxton GD, Sherratt TN, Speed MP, Speed MP, Speed M. 2004 Avoiding attack: the evolutionary ecology of crypsis, warning signals and mimicry. Oxford University Press: Oxford.

6. Stuart-Fox DM, Moussalli A, Marshall NJ, Owens IP. 2003 Conspicuous males suffer higher predation risk: visual modelling and experimental evidence from lizards. Anim. Behav. 66, 541–550.

7. Whiteley AR, Gende SM, Gharrett AJ, Tallmon DA. 2009. Background matching and color-change plasticity in colonizing freshwater sculpin populations following rapid deglaciation. Evolution: Int. J. Org. Evol. 63, 1519–1529.

8. Wente WH, Phillips JB. 2003 Fixed green and brown color morphs and a novel color-changing morph of the Pacific tree frog *Hyla regilla*. Am. Nat. 162, 461–473.

9. Storfer A, Cross J, Rush V, Caruso J. 1999. Adaptive coloration and gene flow as a constraint to local adaptation in the streamside salamander, *Ambystoma barbouri*. Evolution 53, 889–898.

10. McGaugh SE. 2008. Color variation among habitat types in the spiny softshell turtles (Trionychidae: Apalone) of Cuatrociénegas, Coahuila, Mexico. J. Herpetol. 42, 347–354.

11. Rosenblum EB, Römpler H, Schöneberg T, Hoekstra HE. 2010 Molecular and functional basis of phenotypic convergence in white lizards at White Sands. P. Natl. Acad. Sci. USA 107, 2113–2117.

12. Clusella-Trullas S, Wyk JH, Spotila JR. 2009 Thermal benefits of melanism in cordylid lizards: a theoretical and field test. Ecology 90, 2297–2312.

13. van Bergen E, Beldade P. 2019. Seasonal plasticity in anti-predatory strategies: matching of color and color preference for effective crypsis. Evol. Lett. 3, 313–320.

14. Zimova M, Mills LS, Nowak JJ. 2016 High fitness costs of climate change-induced camouflage mismatch. Ecol. Lett. 19, 299–307.18.

15. Olsson M, Stuart-Fox D, Ballen C. 2013 Genetics and evolution of colour patterns in reptiles. Semin. Cell Dev. Biol. 24, 529–541.

16. McLean CA, Stuart-Fox D. 2014 Geographic variation in animal colour polymorphisms and its role in speciation. Biol. Rev. 89, 860–873.

17. Pellitteri-Rosa D, Martín J, López P, Bellati A, Sacchi R, Fasola M, Galeotti P. 2014. Chemical polymorphism in male femoral gland secretions matches polymorphic coloration in common wall lizards (*Podarcis muralis*). Chemoecology 24, 67–78.

18. Cuervo JJ, Belliure J. 2013 Exploring the function of red colouration in female spiny-footed lizards (*Acanthodactylus erythrurus*): patterns of seasonal colour change. Amphibia-Reptilia 34, 525–538.

19. Cadena V, Rankin K, Smith KR, Endler JA, Stuart-Fox D. 2017. Temperature-induced colour change varies seasonally in bearded dragon lizards. Biol. J. Linn. Soc. 123, 422–430.

20. Martín J, Amo L, López P. 2008 Parasites and health affect multiple sexual signals in male common wall lizards, *Podarcis muralis*. Naturwissenschaften 95, 293–300.

21. Sacchi R, Scali S, Pellitteri-Rosa D, Pupin F, Gentilli A, Tettamanti S, Cavigioli L, Racina L, Maiocchi V, Galeotti P, Fasola M. 2010 Photographic identification in reptiles: a matter of scales. Amphibia-Reptilia 31, 489–502.

22. Bergman TJ, Beehner JC. 2008. A simple method for measuring colour in wild animals: validation and use on chest patch colour in geladas (*Theropithecus gelada*). Biol. J. Linn. Soc. 94, 231–240.

23. Sacchi R, Pellitteri-Rosa D, Bellati A, Di Paoli A, Ghitti M, Scali S, Galeotti P, Fasola M. 2013 Colour variation in the polymorphic common wall lizard (*Podarcis muralis*): an analysis using the RGB colour system. Zool. Anz. 252, 431–439.

24. Capolongo D. 1984 Note sull’erpetofauna pugliese. Atti Soc. Ital. Sci. Nat. Mus. Civ. St. Nat. Milano 125, 189–200.

25. Merilaita S, Scott-Samuel NE, Cuthill IC. 2017. How camouflage works. Philos. Trans. R. Soc. B 372, 20160341.

26. West-Eberhard MJ. 2003 Developmental plasticity and evolution. Oxford University Press: Oxford.

27. Michalis C, Scott-Samuel NE, Gibson DP, Cuthill IC. 2017. Optimal background matching camouflage. Proc. R. Soc. B 284, 20170709.

28. Molnár O, Bajer K, Török J, Herczeg G. 2012. Individual quality and nuptial throat colour in male European green lizards. J. Zool. 287, 233–239.

29. Martin M, Meylan S, Gomez D, Le Galliard JF. 2013. Ultraviolet and carotenoid-based coloration in the viviparous lizard *Zootoca vivipara* (Squamata: Lacertidae) in relation to age, sex, and morphology. Biol. J. Linn. Soc. 110, 128–141.

