## Supplementary material, S1 for "Lizard colour plasticity tracks background seasonal changes"

### Supplementary materials – S1

**Some examples of lizards photographically recaptured during field sampling in 2018**

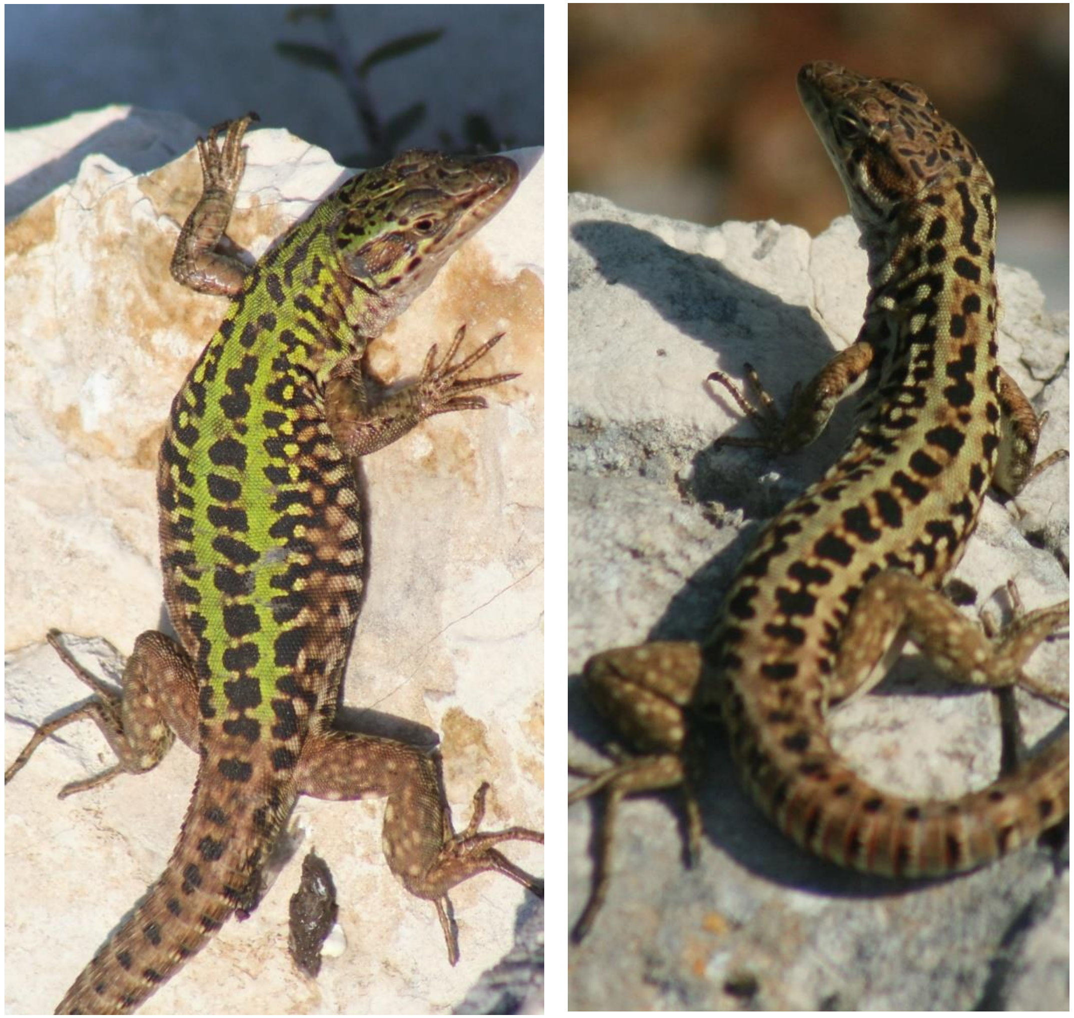

**Figure s1.1** Male photographically captured in March (left) and recaptured in June 2018 (right).

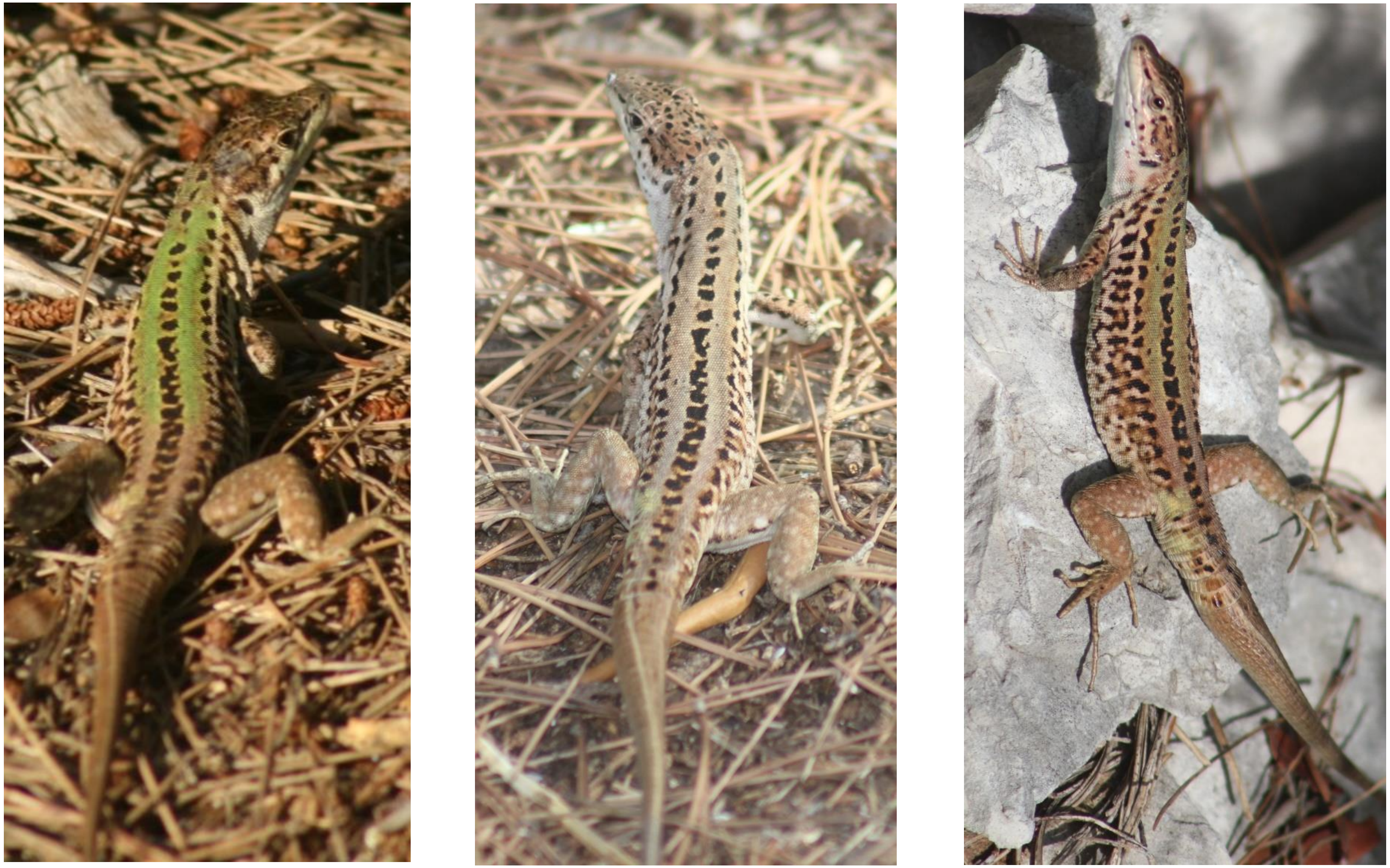

**Figure s1.2** Male photographically captured in March (left) and recaptured in August (middle) and October 2018 (right).

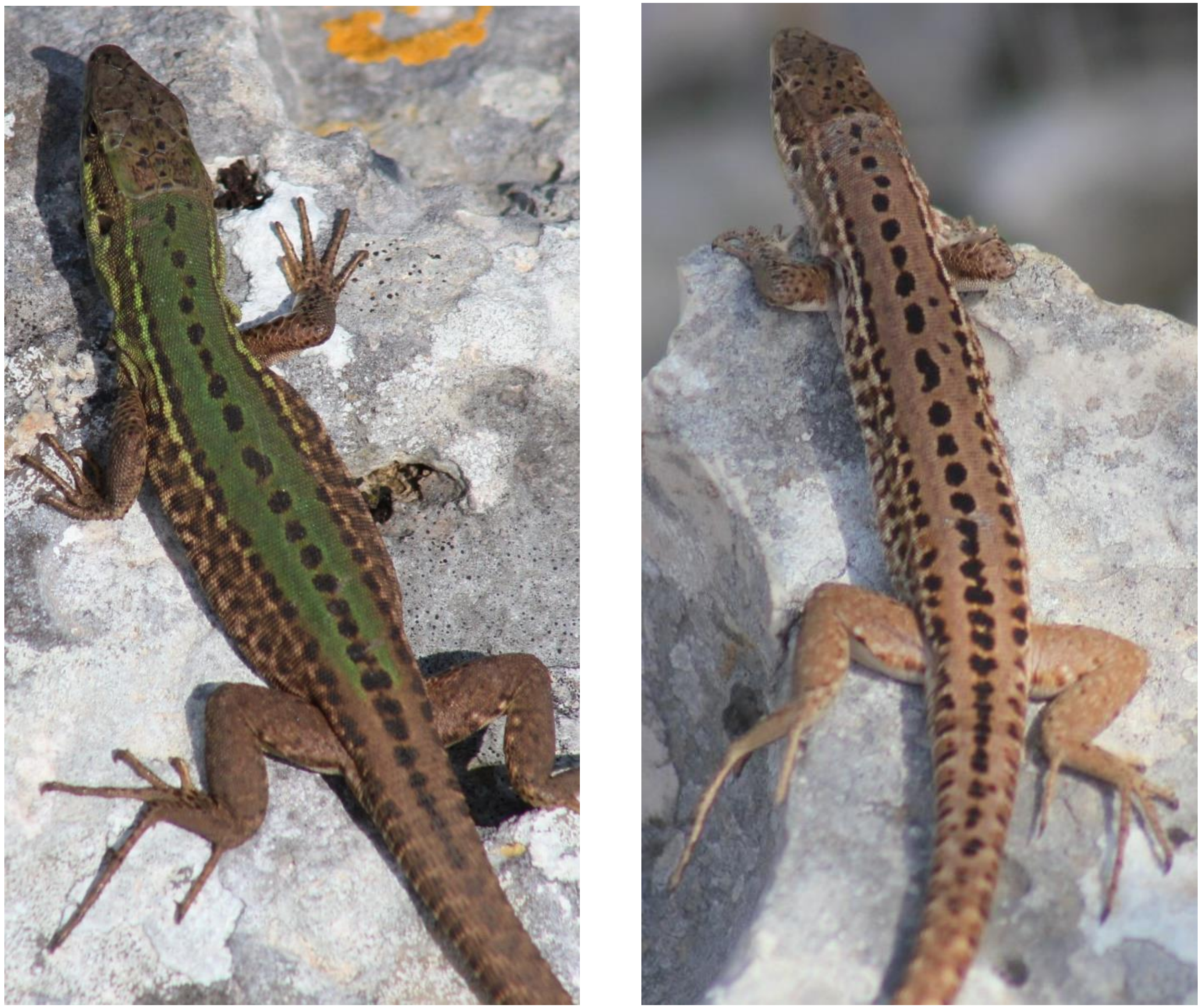

**Figure s1.3** Female captured in March (left) and recaptured in August 2018 (right).

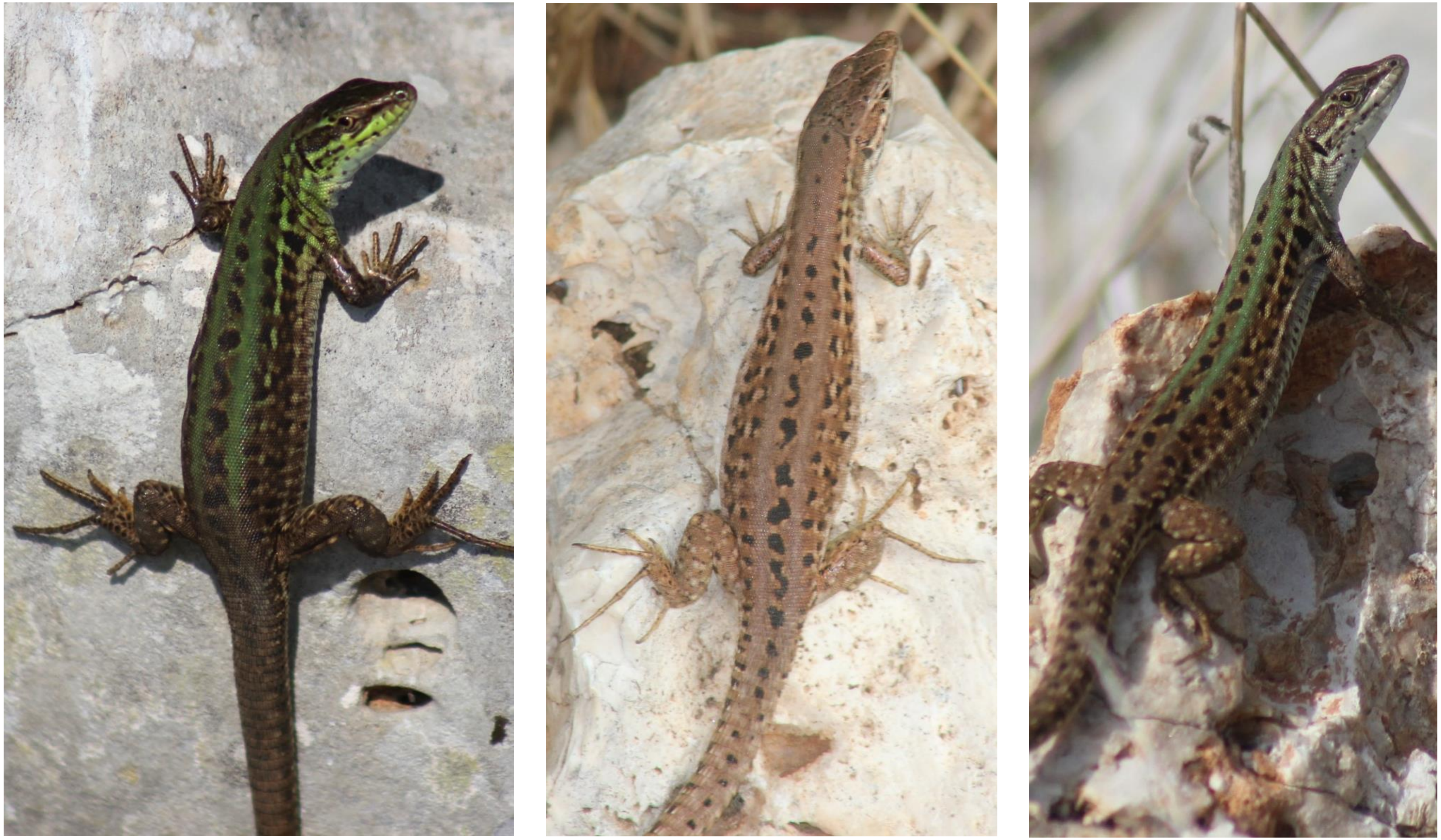

**Figure s1.4** Female photographically captured in March (left) and recaptured in August (middle) and October 2018 (right).
