## Supplementary material, S2 for "Lizard colour plasticity tracks background seasonal changes"

### Supplementary materials – S2

#### Lizards recaptured during field sampling in 2019

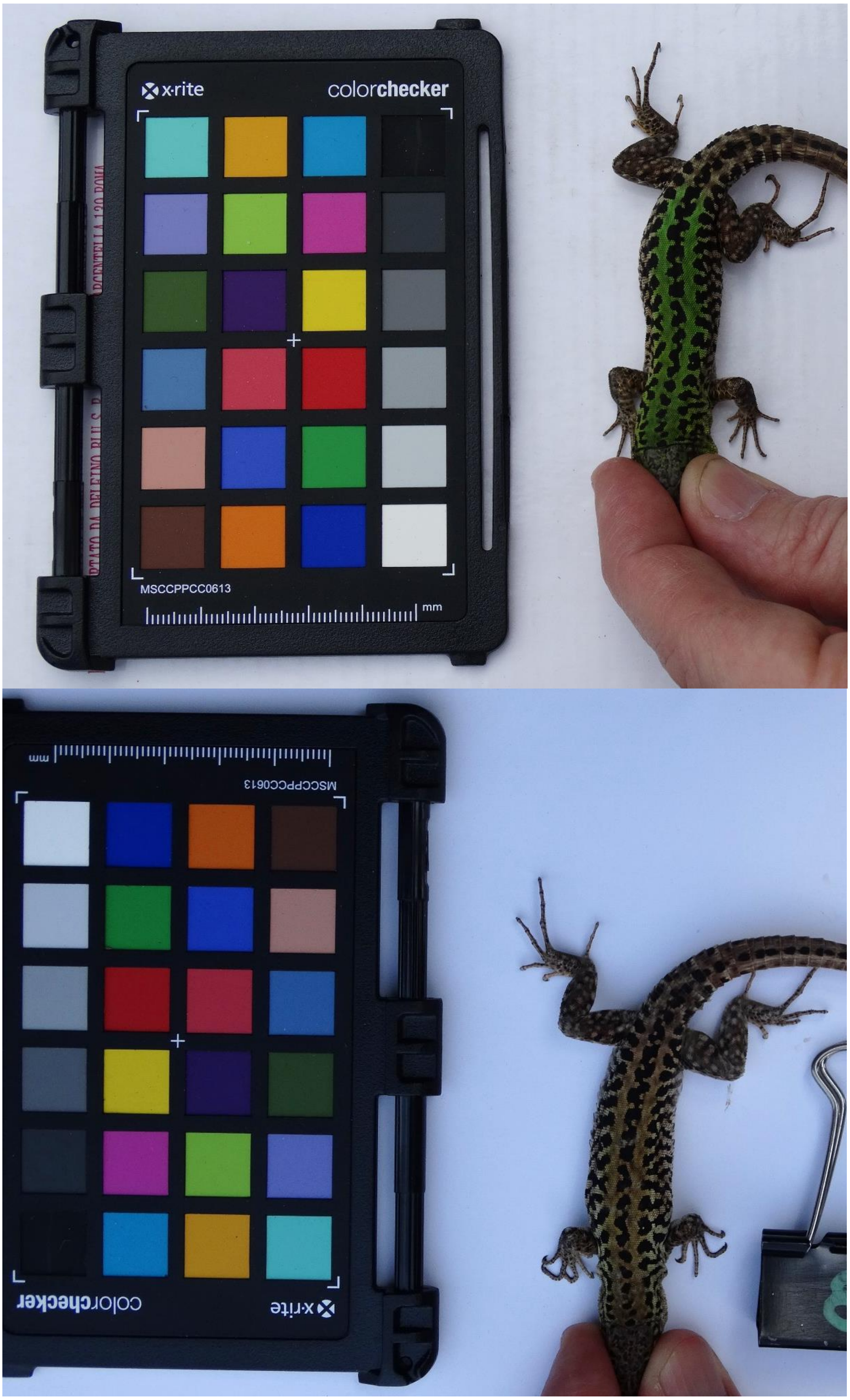


**Figure s2.1** Male captured in March (up) and recaptured in July 2019 (below).

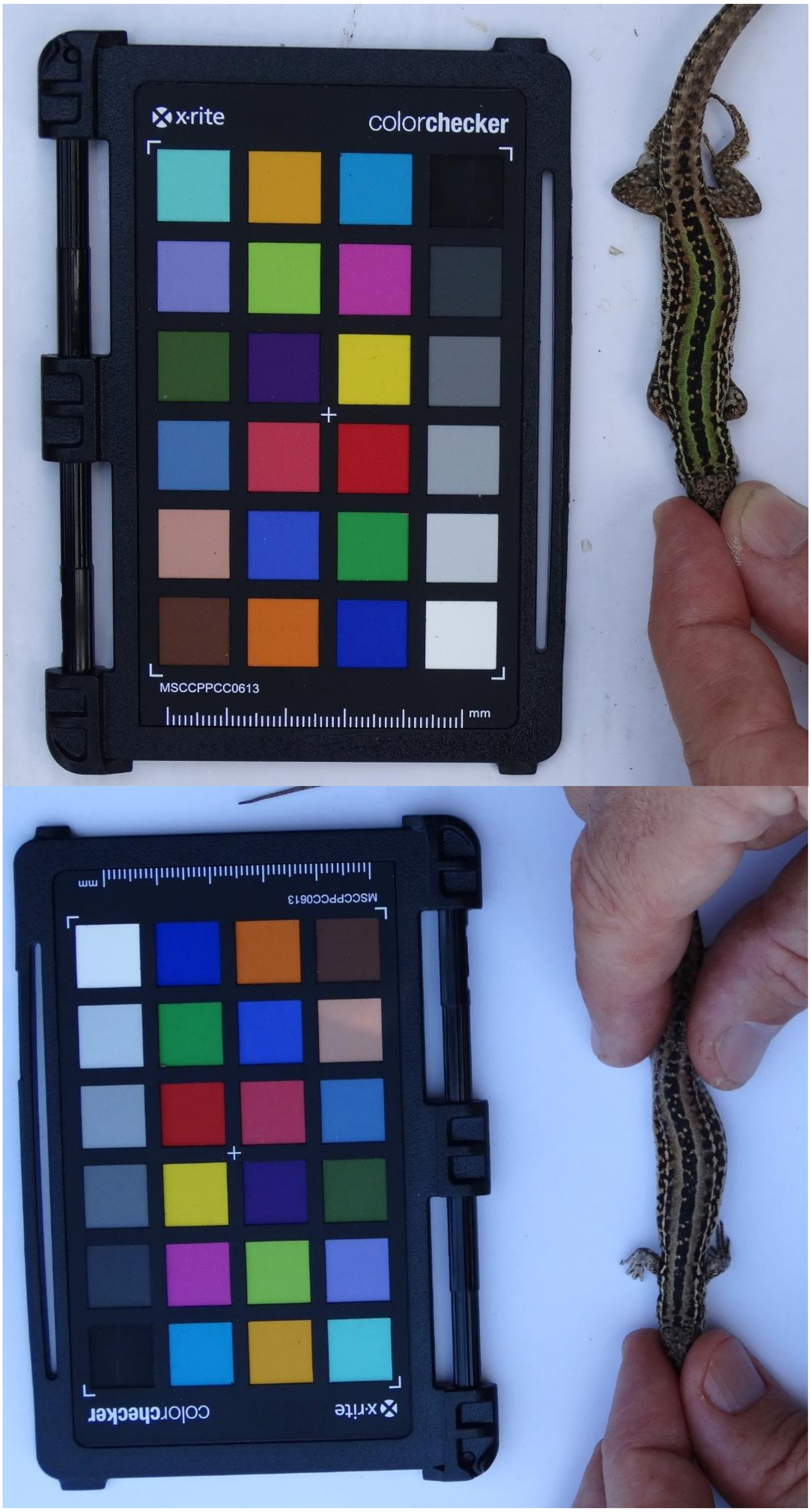


**Figure s2.2** Female captured in March (up) and recaptured in July 2019 (below).

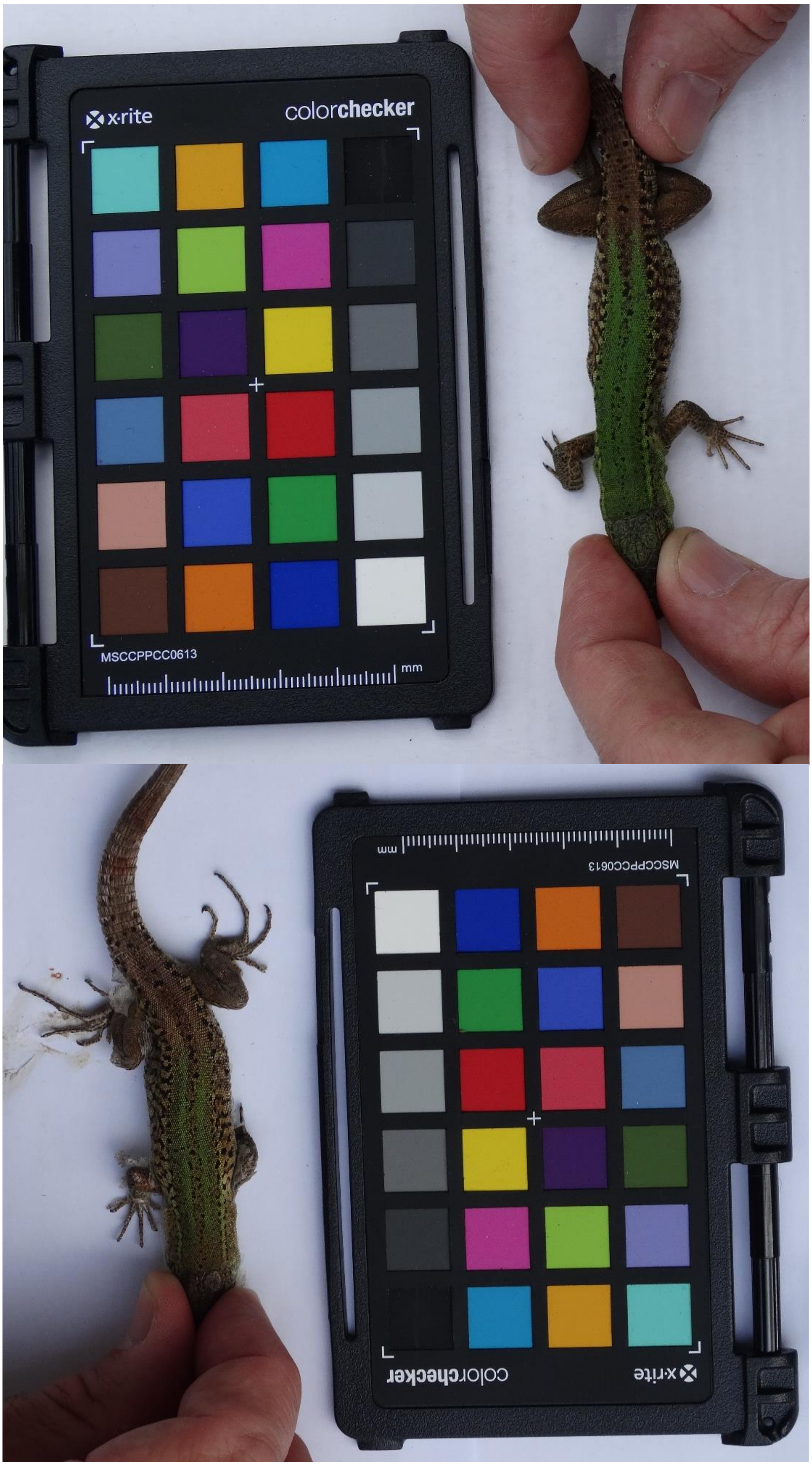


**Figure s2.3** Male captured in March (up) and recaptured in October 2019 (below).

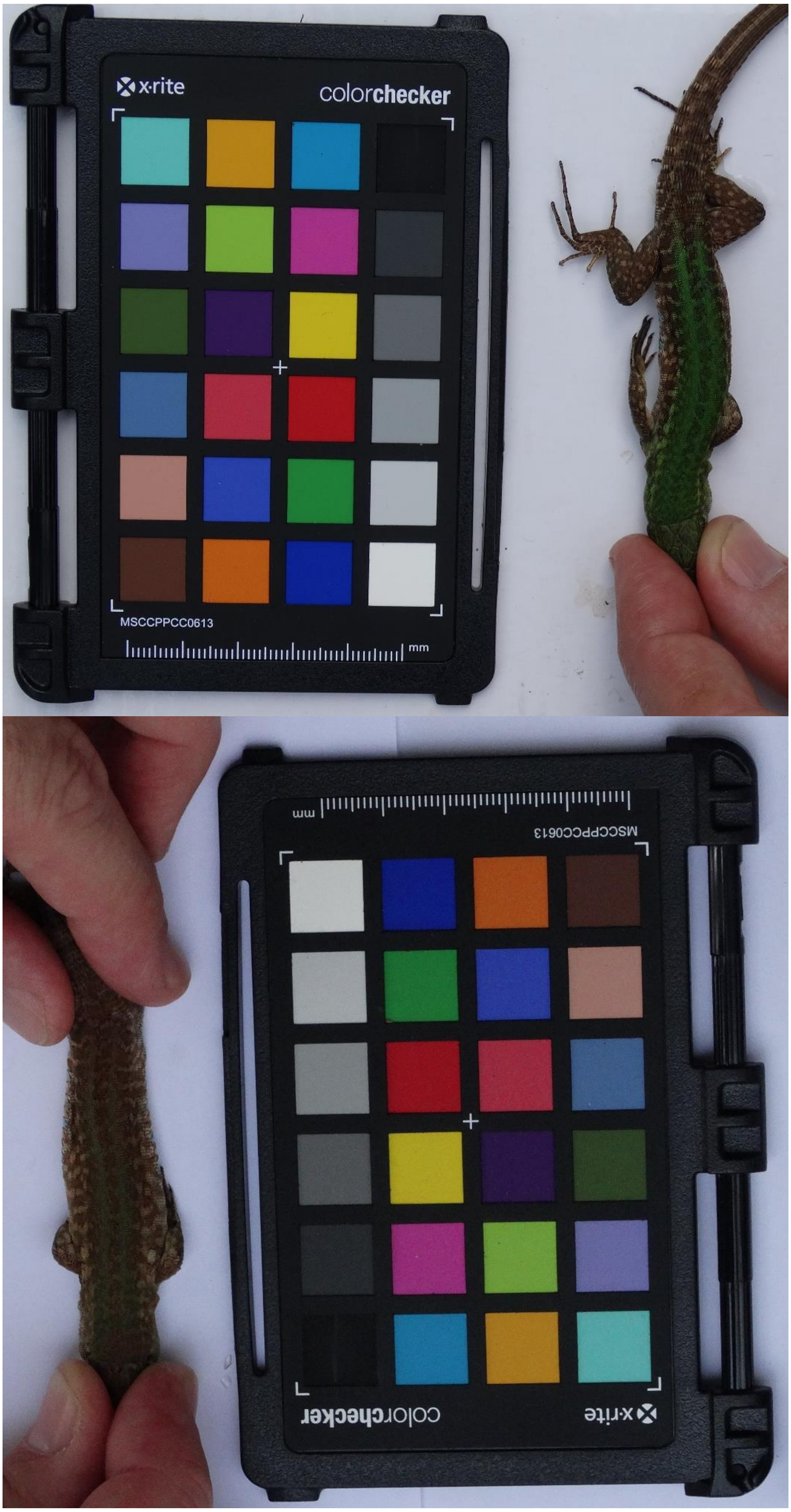


**Figure s2.4** Male captured in March (up) and recaptured in October 2019 (below).

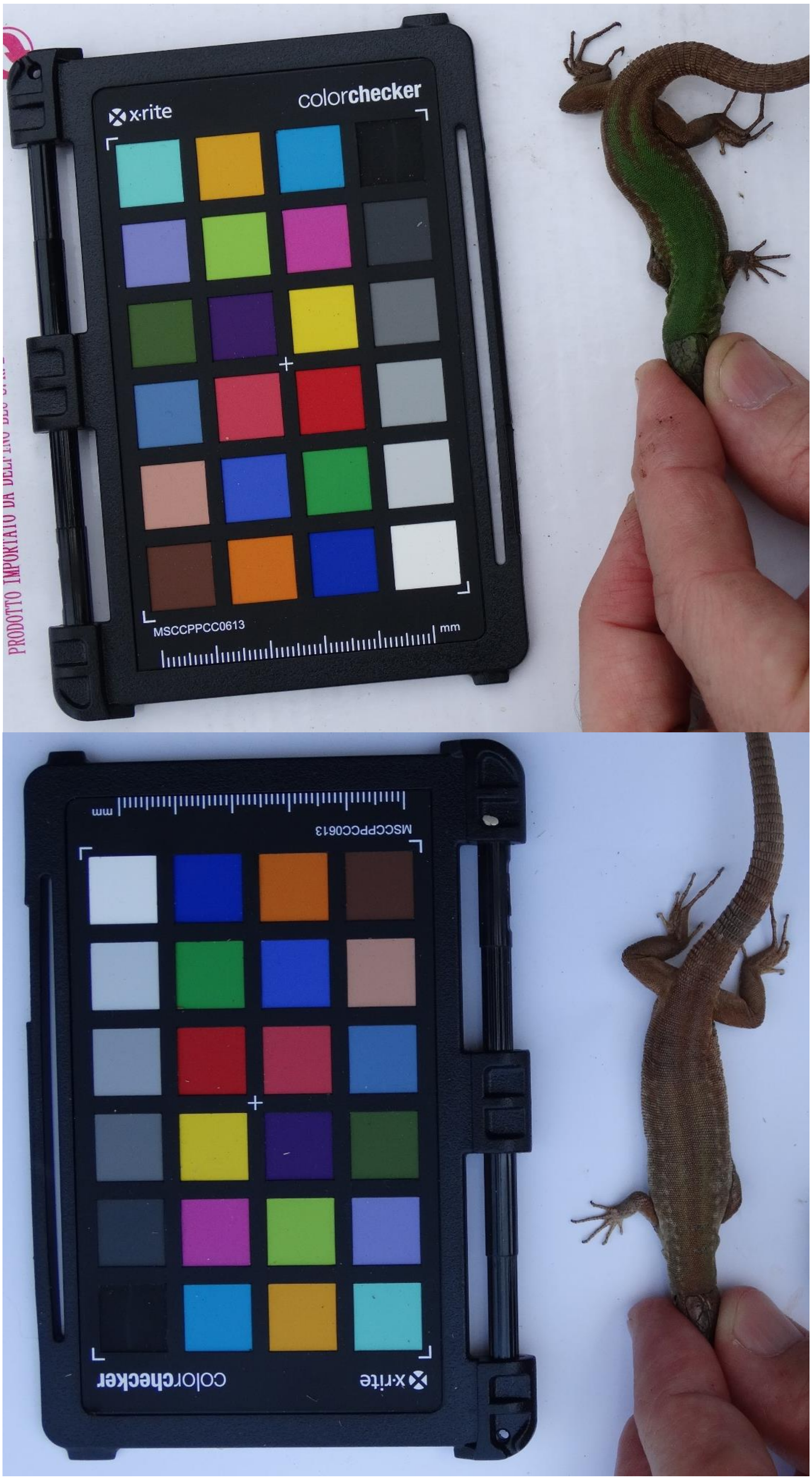


**Figure s2.5** Female captured in March (up) and recaptured in July 2019 (below).
