## Supplementary material, S3 for "Lizard colour plasticity tracks background seasonal changes"

### Supplementary materials – S3

**Examples of background photographed adjacent to the Mini Color Checker chart during 2019**

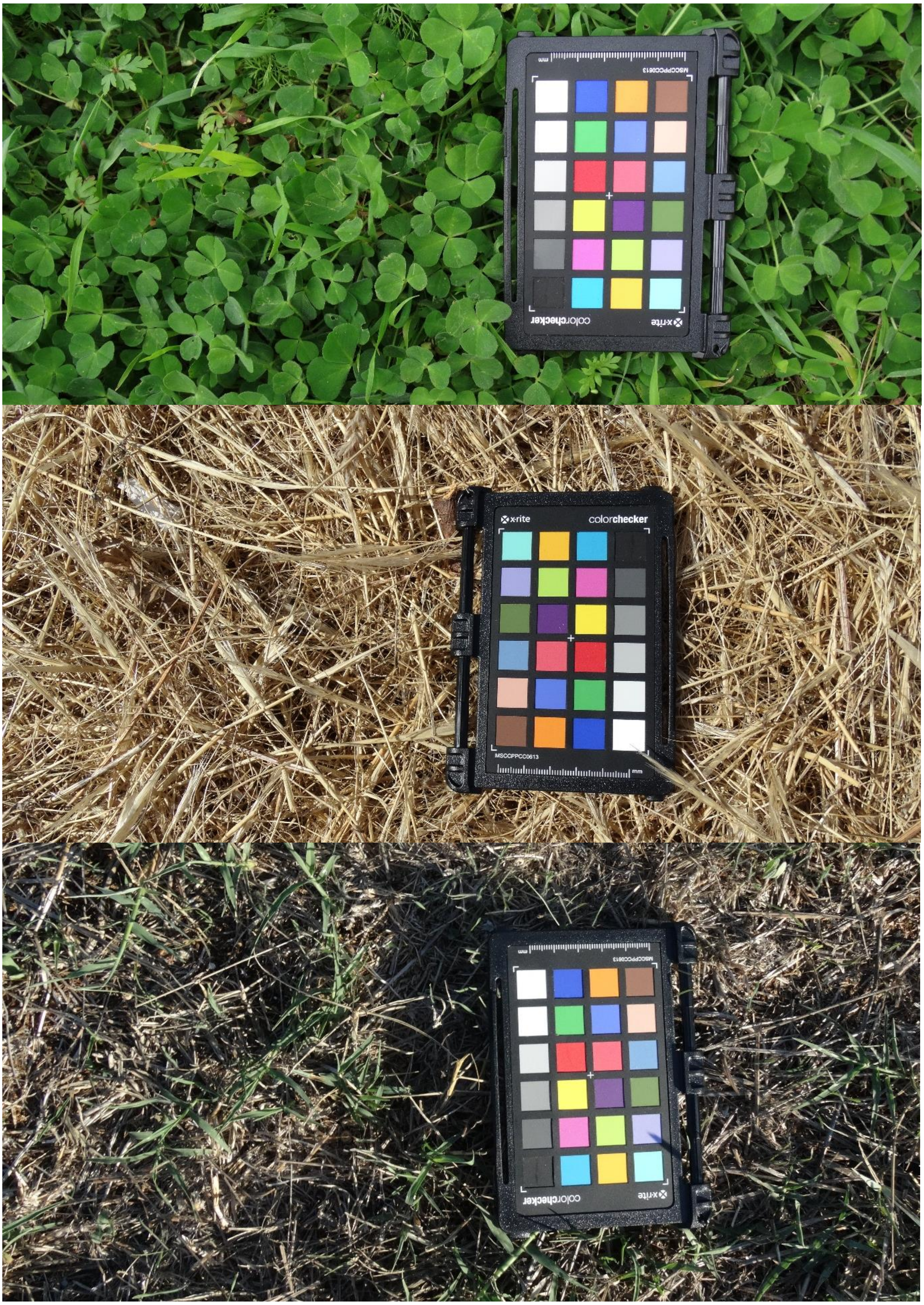


**Figure s3.** Grassy vegetation photographed in March (up), July (middle) and October (below) 2019.
