## Supplementary material, S4 for "Lizard colour plasticity tracks background seasonal changes"

### Supplementary materials – S4

#### Crypsis analysis

When no information on the visual system of potential predators is available, the difference between animal and background colours can provide some information on prey degree of crypsis. Colour dissimilarity can be measured as the Euclidean distance between colours components with a simple formula $D = \sqrt{(H_a - H_b)^2 + (S_a - S_b)^2 + (V_a - V_b)^2}$ [1]. Subscripts a and b refer to animal and background, respectively. In our study, the mean background colour components were calculated from a sample of 20 pictures collected in each different month (March, July, October), and in the same location. In the formula, we inserted the mean value of each background component and the value of each lizard dorsal component, and one distance estimate was calculated for each month.

#### References

1. Endler JA. 1990 On the measurement and classification of colour in studies of animal colour patterns. *Biol. J. Linn. Soc.* **41**, 315–352.

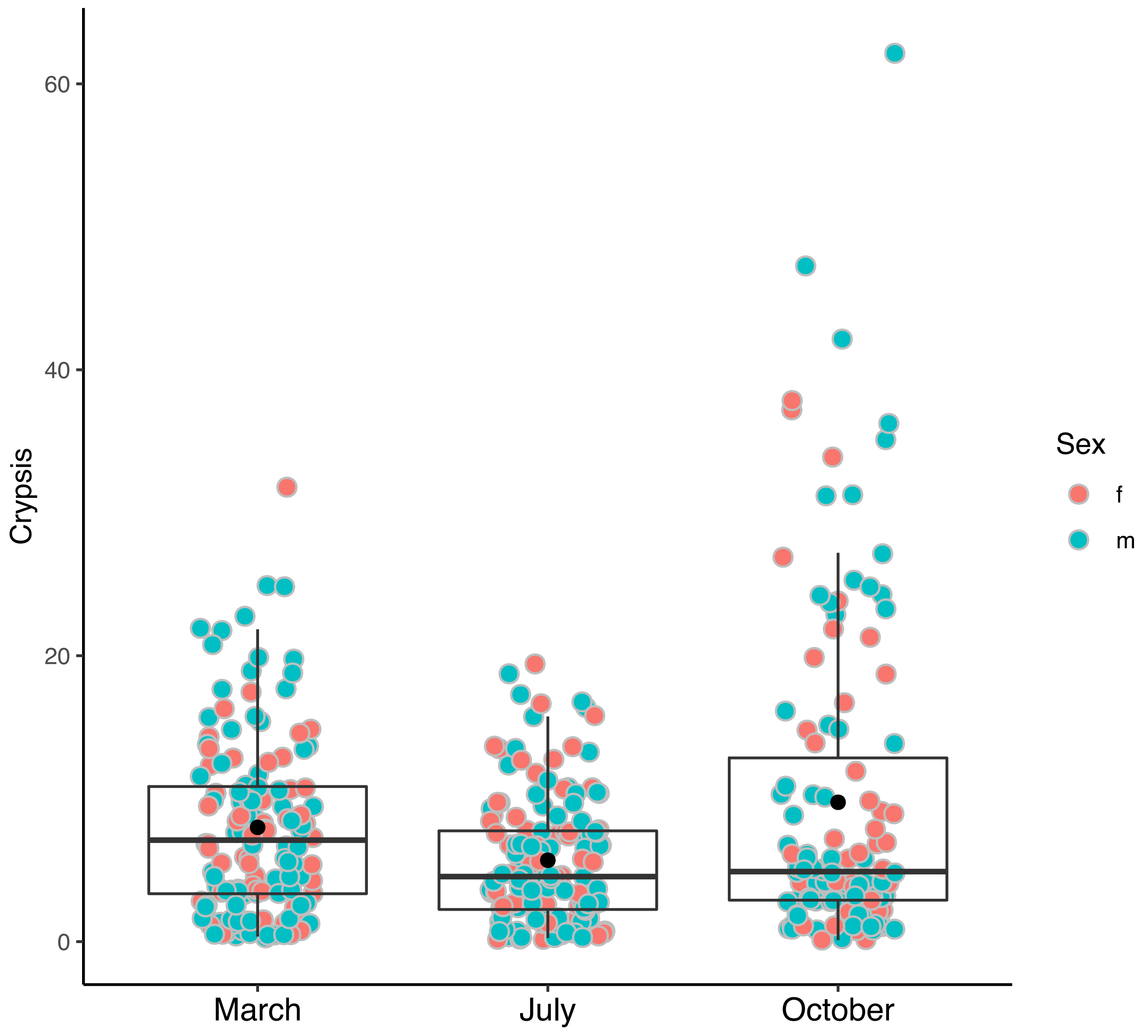


**Figure s4.** Degree of crypsis for each month is reported. Filled black points represent mean crypsis and have been estimated including both sexes.
